# Critical reproductive behaviors in Scaled Quail and Northern Bobwhite are affected by thermal variability and mean temperature

**DOI:** 10.1101/2023.10.20.563339

**Authors:** William Kirkpatrick, Erin Sauer, Rachel Carroll, Jeremy Cohen, Craig Davis, Samuel Fuhlendorf, Sarah DuRant

## Abstract

Animals can respond differently to shifting thermal variability versus thermal averages, both of which are changing due to climate warming. How these thermal variables affect parental care behaviors can reveal the ability of parents to modify their behaviors to meet the competing demands of their offspring’s thermal needs and self-maintenance, which becomes critical in suboptimal thermal conditions. Further, the time frame used to examine the interplay between temperature and behavioral shifts (e.g., seasonal patterns in care vs. drivers of individual care decisions) can provide different information about the plasticity of parental care behavior. We investigated the relationship between thermal means, thermal variability, and incubation behaviors across multiple timescales in Scaled Quail and Northern Bobwhite. Both species decreased off-bout length during periods of high thermal variability, a novel finding among studies of avian parental behavior. Further relationships between thermal endpoints (mean vs. variation) and behavior differed depending on the temporal scale. For instance, total daily time spent off the nest was not influenced by daily average temperature, yet individual off-bout duration increased with increasing average temperature in the two hours prior to the off-bout. These results provide evidence that thermal-behavioral relationships differ across scales and likely represent a bird’s ability to modify their incubation strategy to rapidly respond to the immediate thermal environment (altering individual off-bout length based on temperature) to meet self-maintenance needs while resulting in a similar outcome for their nest (total daily off-bout time). However, longer off-bout durations during high temperature events can come with reproductive costs, sometimes resulting in acute offspring mortality when eggs or chicks experience lethal temperatures.

## Introduction

Moderate projections of anthropogenic climate warming estimate an average global temperature increase of 1.1 – 2.6°C by 2100 (IPCC, 2023) with projected losses of 7.9% of all species due to climate change (Urban, 2015). However, mean increases in temperature alone will not reflect the shifts in thermal conditions that organisms will experience (Denny, 2017; Jensen, 1906; Ruel and Ayres, 1999). Increases in thermal variability are occurring alongside warming average temperatures (Bathiany et al., 2018; Easterling et al., 2000), and evidence in ectotherms shows that animals modify their behavior when exposed to increased environmental variation but not average conditions (Kirkpatrick and Sheldon, 2022; Sheldon and Dillon, 2016; Vasseur et al., 2014). The nature of the relationship between organismal behavior and environmental temperature across genera is complex and often nonlinear (Conway and Martin, 2000), requiring careful contextualization with consideration of multiple aspects of the thermal environment.

When wildlife are exposed to novel thermal conditions, plastic behaviors improve fitness outcomes. This plasticity can have multiple layers of fitness benefits or consequences when the responding behavior is related to parental care, because parental care behaviors have implications for parental and offspring fitness, as well as offspring phenotypic development and plasticity. Behaviors employed by parents to care for their offspring (i.e., parental care behaviors) are known to be plastic (Andes et al., 2020; Dorset et al., 2017) and play an important role in shaping the thermal environment of developing offspring (DuRant et al., 2019). Ambient thermal conditions can alter parental care behaviors (Grimaudo et al., 2020; Masero et al., 2012), subsequently modifying the thermal environment of the developing offspring to directly influence offspring phenotypes (DuRant et al., 2013; Hepp et al., 2006). Examining shifts in parental care behaviors of wild organisms in response to the natural thermal environment can reveal the ability of parents to modify their behaviors to meet the competing demands of offspring care and self-maintenance (McDiarmid et al., 2018; Nelson et al., 2022; Nord et al., 2010; Sharpe et al., 2021; Simeone et al., 2004; Sultan, 2015).

Birds in hot environments must invest in self-maintenance behaviors such as gular fluttering or retreating to cooler microclimates to avoid physiological stress (AlRashidi et al., 2010; van de Ven et al., 2019). However, coping behaviors that cause the parent to leave the nest can come at the cost of offspring care, as the nest is left exposed to external thermal conditions (Carroll et al., 2018; DuRant et al., 2019, 2013; Sharpe et al., 2019). The timing of off-bouts (the time a parent spends off the nest) and their duration play important roles in the development of avian offspring (Cooper and Voss, 2013; Smith et al., 2018). Lengthening and, in some cases, delaying off-bouts can lead to higher rates of predation (Conway and Martin, 2000) or thermal stress, reducing adult and offspring fitness (Nord and Nilsson, 2016; Sharpe et al., 2022).We know little of the scale at which these decisions are being made and their relationship to natural thermal variation (Boulton et al., 2010; Deeming, 2002), but the importance of parental care on offspring development suggests that reproductive success would benefit from the ability of parents to delay off-bouts until temperatures are outside lethal levels for embryos (Smith et al., 2018). Largely, studies focusing on avian behavioral responses to thermal change in temperate climates focus on discrete thermoregulatory behaviors (McDiarmid et al., 2018; Nelson et al., 2022; Walsberg, 1993) or phenological adjustments to rising average temperatures associated with seasonal shifts in the thermal environment (Cohen et al., 2021; Harrod and Rolland, 2020) (McDiarmid et al., 2018; Nelson et al., 2022; Walsberg, 1993). Published data typically show that parents shorten off-bout duration when they are experiencing hot temperatures (Matysioková and Remeš, 2018; Rohwer and Purcell, 2019). The ability of parents to alter their behavior will depend on their ability to withstand physiologically demanding temperatures and to respond to thermal conditions as they are occurring, thus studies examining thermal and behavioral relationships at varying temporal scales are necessary, e.g. acute responses to immediate conditions and broad patterns of behavior across seasonal, monthly or daily scales.

Understanding how the real-time thermal environment (temperatures experienced immediately before exhibition of the observed behavior) influences avian parental behavior is critical for understanding what drives parents to initiate biologically relevant behaviors (Kearney et al., 2012). Long-term (> 24 hours or a single diel cycle) thermal data is useful for exploring seasonal or daily patterns in behavior (Bonamour et al., 2020; Cones and Crowley, 2020; Martin et al., 2018; Simmonds et al., 2017). For instance, greater than average temperature over the course of a reproductive season in birds (i.e., low-resolution thermal and behavioral data) is associated with shorter incubation periods (Capp et al., 2018; Diez-Mendez et al., 2021; Smith et al., 2018), possibly indicating a reduced cost of reproduction for parents. However, assessing only average temperature over the entire incubation stage obscures variance in the thermal environment occurring during this period. During an acute period of hot temperatures, parents may increase nest attendance to shield eggs or seek refuge, leaving eggs exposed to a suboptimal thermal environment. The relationship between thermal variance and incubation behaviors may also vary with scale because thermal variance experienced just before an off-bout may be indicative of thermal stability or rapidly changing temperatures (i.e., rate of temperature change; Sharpe et al., 2021). Temperature variation experienced over a day or entire incubation period, however, may represent the breadth of thermal challenges parents navigated (i.e. the repertoire of behavioral and physiological mechanisms needed to thermoregulate above and below the thermal neutral zone). It is not known how thermal variation shapes incubation behaviors in birds, but acclimation to novel thermal variation and short-term thermal conditions has been evidenced in ectotherms (Fischer and Karl, 2010; Huey et al., 2012; Turriago et al., 2022). What is known about how real-time or recently experienced temperatures shape isolated behavioral responses in birds is mixed, with some species exhibiting reduced off-bout duration in response to increased ambient temperature (Skrade and Dinsmore, 2012; Zhang et al., 2017) and others showing the opposite relationship (Bourne et al., 2021; Caldwell and Cornwell, 1975; Croston et al., 2020; Ringelman et al., 1982). Exploring the effects of acute thermal experiences on variation in avian behavior can inform the mechanisms behind large scale changes in species’ ranges, phenotypic changes, and fitness (Nowakowski et al., 2018).

Here, we describe the variation in the length and timing of off-bout behaviors in response to local thermal conditions experienced throughout the day or immediately prior to the off-bout for two species: Scaled and Northern Bobwhite Quail. We opportunistically used existing data collected in 2015 and 2016 and described in Carroll *et al*. 2018, to begin exploring if both average temperatures and thermal variability experienced by parents shape incubation behaviors. Scaled Quail occupy arid shrublands and prairies from northern Mexico to southern Colorado, USA (Dabbert et al., 2020) while Bobwhite Quail naturally occur in temperate open or early-succession forests and prairies in the eastern USA and Mexico (Brennan et al., 2020). Both of these precocial New World Quail species (family – Ontophoridae) are female-only incubators, but males have been observed in multiple species of the family assisting late in the incubation period (Carroll, 1994). Scaled Quail (*Callipepla squamata*) and Bobwhite Quail (*Colinus virginianus*) overlap in their ranges in western Oklahoma, USA. At range edges, climatic patterns will determine the future of the species in response to climate warming (Rehm et al., 2015), and while the limited overlap in range edges in our species suggests evolutionary distinction, both species have exhibited reduced breeding in response to climatic trends (Carroll et al., 2018; Errington, 1945; Lusk et al., 2006; Pleasant et al., 2006). Nest construction is similar in both species, as both are ground nesters that occupy low-growing trees and shrubs in their shared habitat, though observations suggest that Scaled Quail select nesting locations with more favorable vegetation for thermal buffering than Northern Bobwhite (8.2°C versus 5.7°C cooler nests on average than random locales near the nest, respectively – Carroll et al., 2018).

We hypothesize that: 1) Increased average temperature will lead to increased off-bout duration regardless of temporal scale and species, and 2) thermal variability will operate as a distinct predictor of avian behavior. As mentioned above, little is known about environmental thermal variability as a predictor of behavior, but we expect that parents will likely avoid leaving the nest as often or as long in highly variable conditions to reduce the risk of exposing the nest to temperatures that could quickly become too hot or cold for optimal embryo development. Because parental incubation behaviors can have serious consequences for eggs, we wanted to explore whether temperatures eggs experienced when parents were away from the nest were associated with nest success. Therefore, we also examined nest success (at least 1 egg hatching) in relation to exposure to lethal temperatures (temperatures above 40 °C. We predicted that failed nests would be more likely to experience lethal temperatures and for longer periods of time.

## Methods

### Monitoring Temperature and Incubation Patterns

All nests (37 Bobwhite and 24 Scaled Quail) were monitored in the Beaver River Wildlife Management Area in Oklahoma, USA (36°50′21″N, 100°42′15″W). Environmental temperature data was collected every 30 seconds within and outside the nest using modified Onset Computer HOBOware (Onset Corporation, Borne, MA) thermal sensors. External temperatures were taken with HOBO U23 Pro v2 External Temperature Loggers (Carroll et al., 2018). The internal temperature was taken using egg shaped clay models inserted into the nest that matched the dimensions of quail eggs. Each model egg was inserted into the center of the nest and external sensor was placed within 2m of the nest for standardization of thermal measurements. More details of thermal probes and data collection can be found in Carroll *et al*. 2018. Though thermal-behavioral relationships do change depending on humidity, our field site humidity was not measured at the same frequency or microscale as temperature, so we cannot incorporate it relevantly into our study. According to weather station data collected at a nearby meteorological station (NOAA Station US1OKBV0005 – Forgan, OK, USA), precipitation during the summer was exceedingly rare (and only 9 cm average precipitation on those days).

The machine learning software NestIQ was used to determine off-bout length and timing from the raw nest temperature data (Hawkins and DuRant, 2020). To assess the impact the rate of thermal change has on off-bout duration, we calculated the average and standard deviation of environmental temperature measurements during the two hours before each off-bout. We chose temperatures 2 hours prior to the off-bout to capture a window of thermal temperature long enough to assess a meaningful measure of thermal conditions and stability that could drive an incubation decision without overlapping with longer term diel patterns. To assess the impact the daily thermal environment has on total daily off-bout, we calculated average and standard deviation of temperatures from the morning (sunrise to noon), afternoon (noon to sunset), and night (previous day sunset to sunrise) as well as the temperatures from the previous day and previous afternoon. We included analyses of the temperatures experienced in the previous day, because those conditions could affect incubation decisions by contributing to the parent’s physiological ability to withstand current conditions. We chose these deterministic timeframes to represent a longer-term window of ambient thermal conditions relative to the local diel cycle. We considered including maximum and minimum temperatures from each timeframe, but they both correlated highly with thermal variation (>0.9 and -0.9 respectively) and were excluded. All sunrise and sunset times were recorded for each day of the summers of 2015 and 2016 at the field site and verified through local weather records.

### Statistical Methodology

All statistical analyses were conducted with R version 4.1.2 (R Core Team, 2023). To test the effect of the mean and standard deviation of temperature during the two-hour period before each off-bout on off-bout duration, we developed a linear mixed model (Bates et al., 2014). The universal model included mean and standard deviation of the two-hour period preceding the off-bout as predictors and off-bout length (in minutes) as the response variable. We included a two-level categorical predictor for whether the off-bout took place in the morning or afternoon, as each species typically takes two off-bouts per day, in the early morning or late afternoon (referred to as AMPM in models). We also included species (Scaled Quail or Northern Bobwhite). To control for non-independence of the measurements at a single nest and year, we included individual nest ID, year, and date as random intercepts. All possible interactions between predictors were included in the universal model. To test for relationships between temperature and off-bout start time, we conducted a separate linear mixed model with the same predictors, interactions, and random intercepts described above but with off-bout timing as the response variable (Table 1).

**Table 1.** The best fit models for each suite of thermal predictors as selected by AICc using the package MuMIn. High-resolution: The universal model included mean and standard deviation of the two-hour period preceding the off-bout as predictors and off-bout length (in minutes) as the response variable. We included a two-level categorical predictor for whether the off-bout took place in the morning or afternoon (referred to as AMPM) and species (Scaled Quail or Northern Bobwhite). Low-resolution: All models were structured as above except with averaged thermal data over 6 deterministic timeframes as predictors: 24hr (temperatures from 12AM-11:59PM of the current day), Morning (sunrise to 12PM), Afternoon (12PM to sunset), Previous Night (sunset to sunrise of previous night), Previous Day (12AM to 11:59PM of the previous day), and Previous Afternoon (12PM to sunset of the previous day)). We included individual nest ID, year, and date as random intercepts in all models. All possible interactions between predictors were included in each universal model.

| High-Resolution Best Fit Models |  |  |  |  |  |  |
| --- | --- | --- | --- | --- | --- | --- |
| Model Name | Response Variable | Main Effect(s) | Interactive Effects | AIC<sub>c</sub> | df | AIC<sub>c</sub> w<sub>i</sub> |
| Individual Duration | Off-bout Duration | 2hr Mean, 2 hr SD, AMPM, Species | 2hr Mean: Species, 2hr SD: Species, AMPM: Species | 8940 | 12 | 0.80 |
| Null | Off-bout Duration | ~ | ~ | 9156 | 5 | 1.2e-47 |
| Low-Resolution Best Fit Models |  |  |  |  |  |  |
| Model Name | Response Variable | Main Effect(s) | Interactive Effects | AIC<sub>c</sub> | df | AIC<sub>c</sub> w<sub>i</sub> |
| Total Duration | Daily Off-bout Duration | 24hr SD, Morning SD, Species | 24hr SD: Species, Previous Night SD: Species | 5666 | 10 | 0.02 |
| Null | Daily Off-bout Duration | ~ | ~ | 5706 | 4 | 6.2e-11 |
| Morning Start Time | Off-bout Timing - AM | Previous Day SD, Species | ~ | 1367 | 7 | 0.12 |
| Null | Off-bout Timing - AM | ~ | ~ | 1387 | 5 | 5.8e-0.6 |

To address the effect of temperatures experienced over a longer time period on total daily off-bout, we developed a linear mixed effects model which included the sum duration of all off-bouts in 24 hours as the response variable. We selected total daily off-bouts as our response variable because the predictors are date-specific thermal measurements instead of off-bout specific thermal measurements. The predictive variables included the mean and standard deviation of external temperature from the following timeframes relative to the date: 24hr (temperatures from 12AM-11:59PM of the current day), Morning (sunrise to 12PM), Afternoon (12PM to sunset), Previous Night (sunset to sunrise of previous night), Previous Day (12AM to 11:59PM of the previous day), and Previous Afternoon (12PM to sunset of the previous day). We chose these time periods to allow for comparison of longer-term trends in temperature. While they are likely highly correlated with one another, our model selection strategy (see below – *Model Selection*) corrected for this issue. We included a two-level categorical predictor for species (Scaled or Northern Bobwhite). To control for non-independence of the nest location and year, we included individual Nest ID and year as random intercepts. To test for relationships between mean and standard deviation of temperature on the average daily time of each off-bout, we conducted two separate linear mixed models with the same predictors, interactions, and random intercepts described above, but the response variables were the timing of the morning or afternoon off-bout respectively, given that each species takes at most, two off-bouts during the day in the morning, evening, or both.

### Model Selection

For each universal mixed-effects model (high and low-resolution), we used the dredge function from the MuMIn package for model selection (Bartoń, 2022). We chose the best fit model by finding the model with the lowest relative AICc score (Burnham et al., 2011). All values represented in our figures are partial residual plots generated using the predict function in R.

The best fit model for the high-resolution approach to analyze individual off-bout duration (referred to in Results as Individual Duration) included 2-hour mean temperature, 2-hour standard deviation of temperature, morning or afternoon (AMPM), and species as main effects with three interactive effects: 2-hour mean*species, 2-hour standard deviation of temperature*species, and AMPM*species (Table 1).

The best fit model for predicting total daily off-bout (referred to in Results as Total Duration) included 24 hour and morning standard deviation of temperature as main effects; standard deviation of morning temperatures was removed due to a high correlation with 24 hour variation (r^2^ > 0.6) and lack of a significant effect in the general model. Species and two interactive effects were also included: species*morning standard deviation of temperature and species*previous night standard deviation of temperature (Table 1).

Two models were used to predict daily off-bout start time, one that included morning off-bouts (referred to in Results as Morning Start Time) and one that included afternoon off-bouts since each species takes two off-bouts per day in either the morning or evening. The best fit model for the afternoon off-bouts (Total PM Start Time) included mean temperature during the afternoon and species with no interactive effects (Table 1).

### Conditional Model Residuals

Behavioral responses to temperature in wildlife are often nonlinear, especially in extreme thermal scenarios (Denny, 2017). By examining the conditional residuals of our fitted models, we can determine linearity and heteroscedasticity of the relationship observed (Santos Nobre and da Motta Singer, 2007). For the model Individual Duration, the residuals were largely linear, with a slight deviation above expected observations as residual values increased (Supplementary Figure 2). However, non-linear and/or heteroscedastic relationships were noted when examining timing of off-bouts on local temporal scales. Models predicting high-resolution off-bout start time and morning off-bouts did not meet the necessary assumptions. Therefore, we only examined models that met the requirements for heteroscedasticity and non-linearity (see Supplementary Materials to reference conditional residual plots).

### Hatching Success

We compared cumulative minutes exposed to temperatures above 40°C in hatched vs. failed nests. We also constructed a similar model with days exposed to temperatures above 40°C to see whether failed nests were exposed more frequently to lethal temperatures. To determine the effect of time exposed to extremely warm conditions, we constructed two one-way ANOVA’s with either total minutes above 40°C experienced or days above 40°C. The models included nest hatching or failure and species as main effects. Interactive effects of species and nest failure were also included.

## Results

### High-Resolution Thermal Data – Duration

As described by model Individual Duration, off-bout duration increased as mean temperature increased (β = 4.64 ± 0.46, t = 10, p < 0.001; Figure 1A). However, in response to increased thermal variation, lengths of individual off-bouts decreased (β = -4.68 ± 2.11, t = -2.2, p = 0.03; Figure 1B).

**Figure 1.**
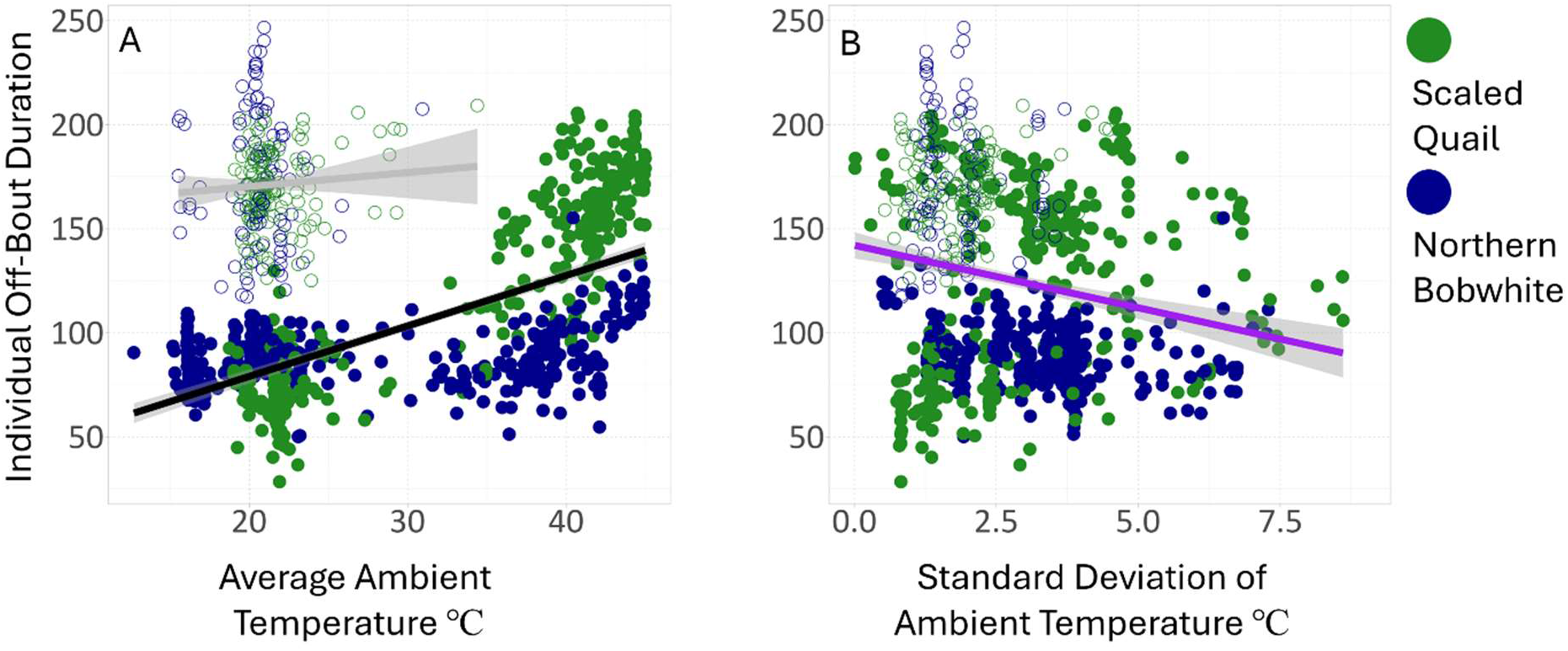
Partial residual plots showing the relationships between the thermal environment immediately preceding individual off-bouts and off-bout duration. Blue lines exhibit the association across both years while black and gray lines represent 2015 and 2016, respectively. Open circles represent quail nests in 2015, and closed circles represent 2016 nests. A) As mean temperature in the 2 hours preceding each off-bout increased, Northern Bobwhite (green) and Scaled Quail (blue) off-bout duration increased. Nest sites in 2015 were cooler on average than 2016 (2015 = 21.5 C, 2016 = 31.2 C). B) As standard deviation of temperature increased, individual off-bout duration decreased. All points are predicted model values.

Northern Bobwhite averaged 140 ± 68 minutes per off-bout while Scaled Quail averaged 110 ± 57 minutes (β = 62 ± 14.7, t = 4.2, p < 0.001). Northern Bobwhite exhibited longer afternoon off-bouts (159 ± 61 minutes) than Scaled Quail (Afternoon =108 ± 48 minutes; β = - 43.1 ± 10.5, t = -4.1, p > 0.001) even when mean temperatures 2 hours preceding the off-bout were high (β = -1.8 ± 0.7, t = -2.7, p = 0.007, Supplementary Figure 2B). For both species, morning off-bouts were not significantly longer than afternoon off-bouts, though they were different (β = 1.5 ± 7.4, t = 0.2, p = 0.8; Morning = 110 ± 66 minutes; Afternoon = 138 ± 61 minutes).

**Figure 2.**
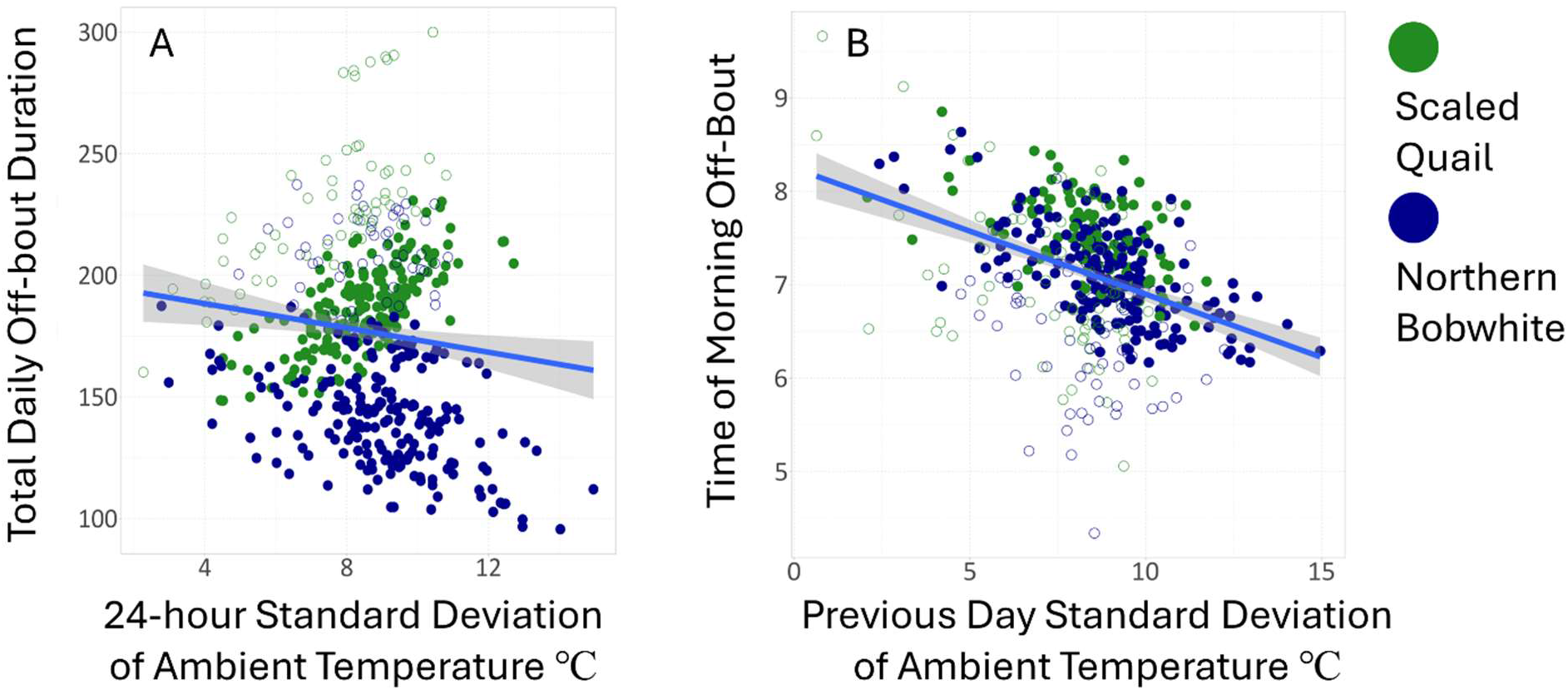
Partial residual plots showing the relationships between the thermal environment measured over selected timeframes, total daily off-bout duration (A), and morning off-bout initiation (start time) (B). A) As thermal variation increased, both species reduced their total daily off-bout duration. B) Thermal variation of air temperature of the day preceding the morning off-bout was a significant negative predictor of start time, advancing off-bout initiation. Open circles represent quail nests in 2015, and closed circles represent 2016 nests.

### Low-Resolution Thermal Data – Total Daily Off-Bout

As described by model Total Duration, as thermal variability of the total day (24 hour standard deviation) increased, daily off-bouts lengthened (β = 11.9 ± 3.7, t = 3.2, p = 0.001; Figure 2A). Alternatively, thermal variation of the previous night was selected for the model but was not a significant predictor of total daily off-bout (β = 5.6 ± 6.4, t = 0.9, p = 0.38). Scaled Quail spent less total time off the nest daily (156 ± 74 minutes) than Northern Bobwhite (197.2 ± 81 minutes; β = 71 ± 34, t = 2.1, p = 0.03).

### Low-Resolution Thermal Data – Start Time

As described by model Morning Start Time, increasing thermal variation of the previous day advanced the initiation of the following day’s off-bouts into the cooler early morning (β = - 0.16 ± 0.03, t = -4.9, p < 0.001; Figure 2B)

### Hatching Success

Both species constructed nests in locations that experienced temperatures above the critical thermal maxima for eggs (∼40°C and above). Our analysis revealed a marginal difference in total minutes exposed to 40°C temperatures in failed vs. hatched nests (one-way ANOVA; *df* = 1,31, *F* = 4.1, *p* = 0.051). For both species, failed nests experienced 40°C+ for 98 ± 133 (mean ± SD) minutes compared to 30 ± 56 minutes in successful nests. There was not a significant effect of the number of days exposed to temperatures > 40°C or species, though failed Northern Bobwhite nests were typically exposed to less time above the threshold than Scaled Quail nests (88 minutes ± 140 and 111 ± 135 minutes respectively).

## Discussion

We investigated the relationship between average thermal conditions, thermal variability, and incubation behaviors across multiple timescales in Scaled Quail and Northern Bobwhite. We provide evidence that both species reduced time on the nest following short periods of increased thermal variability (i.e.at high rates of temperature change), a novel finding among studies of avian parental behavior. Further, relationships between thermal endpoints (mean vs. variation) and behavior differed depending on scale, in other words, whether we examined environmental temperature just prior to an individual off-bout or thermal conditions across the course of a day and daily incubation patterns. These results provide evidence that the thermal factors shaping discrete behavioral decisions may be washed out when assessing general patterns of behavior over longer time scales (Garcia et al. 2019) and both are important to consider when estimating effects of climate change on reproductive patterns.

Northern Bobwhite and Scaled Quail exhibited plastic behavioral responses to recently experienced temperatures. Both species shortened off-bouts in response to increasing thermal variability (i.e. higher rate of temperature change; Figure 1B). Similarly, total time away from the nest during a 24 hour period decreased with increasing daily thermal variability (Figure 2A, B). Alternatively, increases in recently experienced average temperatures caused parents to increase the length of an individual off-bout (Figure 1A), but daily average temperatures were not important to the total time spent away from the nest in a 24 hour period. These findings demonstrate that thermal variability can act independently of average temperature to shape an important reproductive behavior in birds and the directionality of these relationships can vary with scale. Presumably, birds shortened the length of an individual off-bout with increasing thermal variability because temperature was rapidly changing or unpredictable and could quickly reach lethal temperatures or temperatures that would slow embryonic development. Our study cannot distinguish between decisions driven by parental self-maintenance needs and offspring requirements, so shorter off-bouts during high rates of temperature change could also benefit parents. It is less clear why daily thermal variability would shorten total time away from the nest that day, but it may be that temperatures were changing most dramatically at the times of day that the quail would typically take an off-bout, shortly after sunrise and late afternoon. Alternatively, high daily thermal variability could generally signal that temperatures are unpredictable, causing parents to shorten all incubation breaks that day. The increased care we detected in our study is similar to findings in smallmouth bass (*Micropterus dolomieu*). Bass nesting in more variable habitats increased parental care activities compared to parents nesting in less variable but warmer habitats (Cooke et al., 2003). It was surprising that relationships between daily patterns of incubation and temperature did not match the relationship between off-bout length and the average temperature experienced just before the off-bout. This suggests that temperatures experienced at the time of an off-bout do inform incubation decisions, and these patterns will not always scale up to be reflected in daily or seasonal patterns in incubation constancy (Grubbauer and Hoi, 1996; Husby et al., 2010; Le Vaillant et al., 2021; MacDonald et al., 2014; Taff and Freeman-Gallant, 2021; Zhang et al., 2017). This finding is supported by studies of other species nesting in hot conditions. Least Tern and Piping Plover (*Charadrius melodus*; Andes et al., 2020) incubating parents exhibit some behaviors during incubation that aid in their thermoregulation but only within certain ranges of environmental temperature and these relationships would not be obvious when summing time away from the nest across a longer time period. These individual incubation decisions based on current thermal conditions can have important implications for nest success. We found that most failed nests were exposed to temperatures exceeding 40°C on a single day in a single extreme event, this finding suggests that a single exposure to extreme heat during an off-bout can lead to reproductive failure (and see Cunningham et al., 2013) and may not be detectable when comparing general patterns of incubation in relation to average temperature. In some species, there is evidence of population collapse caused by a short exposure to extreme heat regardless of corresponding daily mean temperature or parental responses to temperature (Ma et al., 2015).

When comparing behaviors of the two quail species, we found that both species expressed similar trends in their behavioral modifications to thermal conditions but differed in individual off-bout duration and temperature of their nests. Scaled Quail spent significantly less time off the nest on average (156 minutes) than Northern Bobwhite (197 minutes). Species specific differences in these data were explored elsewhere (Carroll et al. 2018), including this particular result, so we discuss it here as it relates to the hypotheses we tested. It is interesting that the two quail species differ in off-bout duration, but not in the relationship between the responsiveness of their incubation behaviors to thermal conditions, both thermal averages and variation. Off-bout duration and nest temperatures are likely influenced by natural history differences of the two species (Brennan et al., 2020; Dabbert et al., 2020) and determined primarily as a result of Scaled Quail selecting nest sites or substrates with superior thermal buffering characteristics compared to Northern Bobwhite nests (Carroll et al., 2018). The typical breeding habitat of Northern Bobwhite is temperate and extreme temperatures rarely reach those experienced by Scaled Quail (Brennan et al., 2020), which are a grassland desert-dwelling species adapted to the extreme highs and lows present in the Western Oklahoma field site (Dabbert et al., 2020). The two species may have experienced different selection pressures for nest site selection, but plasticity of incubation behaviors to changing thermal conditions indicate the importance for both species to prevent exposure of nests to lethal temperatures and reduce care costs to the parent. It should be noted that direct comparisons between each species based on results found here need to be made with caution, as thermal and behavioral measurements across each species’ range would be required to make definitive species level comparisons in behavioral plasticity.

Behavioral plasticity during breeding generally benefits both the parent and dependent offspring by keeping the thermal environment stable over time (Kirkpatrick and Sheldon, 2022; Mamantov and Sheldon, 2021; Telemeco et al., 2009), but shifts in behavior alone cannot completely negate the impacts of poor or unexpected environmental conditions (Sharpe et al., 2019, 2022; Sudnick et al., 2021), as time allotted for self-maintenance may take precedence over offspring care (Deeming, 2002). Our results indicate that the rate of environmental temperature change can directly influence parental behavior, highlighting that temperature variability may be a key player in shaping avian reproductive behaviors. Our analysis also indicated that discrete behavioral decisions (timing or duration of an off-bout) are made using thermal information about current conditions, but more general patterns in incubation behaviors (total daily time spent off the nest) were less responsive to the daily average temperature. In other words, in these two species, parents are spending a similar amount of time off the nest each day regardless of average daily temperature, but how that time is metered out is flexible based on recently experienced thermal conditions and temperature stability. This finding underscores the importance of testing for relationships between thermal conditions and behavior across multiple timescales. A single thermal measurement taken over a defined time may provide inaccurate assessments of organismal vulnerability to thermal change if characterized incorrectly or incompletely (Clusella-Trullas et al., 2021; Sheldon and Dillon, 2016). Models with averaged, lone sole-characteristic predictors should not be avoided but contextualized temporally as we do here (Garcia et al., 2019).

## CRediT Authorship Statement

William Kirkpatrick: Conceptualization, Writing - Original Draft, Writing – Editing and Review Methodology, Data Curation, Formal Analysis. Erin Sauer: Formal Analysis, Writing – Editing and Review. Rachel Carroll: Investigation, Resources, Methodology, Data Curation. Jeremy Cohen: Formal Analysis, Writing – Editing and Review. Craig Davis: Investigation, Resources, Methodology, Data Curation, Funding Acquisition. Sam Fuhlendorf: Investigation, Resources, Methodology, Data Curation, Funding Acquisition. Sarah DuRant: Conceptualization, Investigation, Resources, Methodology, Data Curation, Writing - Original Draft, Writing – Editing and Review, Supervision.

## Acknowledgements

We would like to thank Rachel Carroll, Craig Davis, and Sam Fuhlendorf for permission to use their thermal data from Scaled and Northern Bobwhite Quail nests. Thanks to Wayne Hawkins for assistance with NestIQ.

## Ethical Committee/Institutional Review Board (IRB) Approval

Animal welfare approval was obtained from IACUC at Oklahoma State University.

## Data Accessibility Statement

All supporting data and annotated R scripts are available in <u>GitHub</u> or through contact with the corresponding author.

