## Supplemental Materials for "Critical reproductive behaviors in Scaled Quail and Northern Bobwhite are affected by thermal variability and mean temperature"

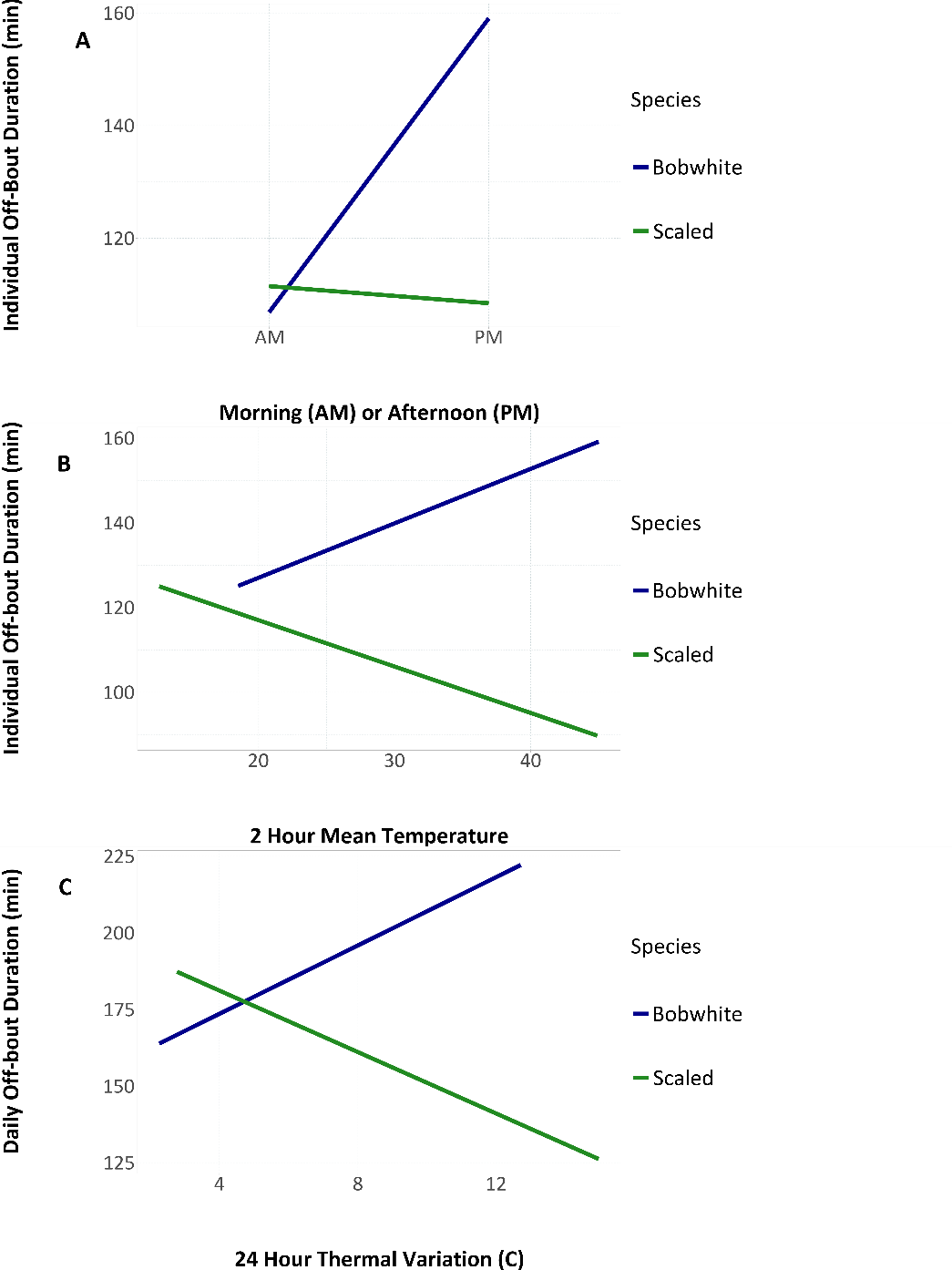
**Supplementary Materials**

Supplementary Figure 1. A) There was a significant interactive effect of whether the off-bout took place in the morning or afternoon; Bobwhite Quail take longer afternoon off-bouts than the desert-acclimated scaled quail (β = -43.1 ± 10.5, t = -4.1, p > 0.001; Bobwhite: 159 ± 61 minutes; Scaled: 108 ± 48 minutes). For the high-resolution (B: β = -1.8 ± 0.66, t = -2.7, p = 0.007) and low-resolution scales (C: β = -9.9 ± 3.8, t = -2.6, p = 0.01), Northern Bobwhite exhibited increased off-bout duration in response to thermal averages and standard deviation, respectively. Scaled Quail, as reported in the main findings, did as well. Despite the significant interactive effects reported above, the reported opposing effect was due to 2015 being significantly cooler, causing the data to have an odd grouping (as evidenced in Figure 1) that suggested negative effects where none existed when individual nest plots were examined. Year was considered as a random effect in all analyses, with residual plots and individual nest
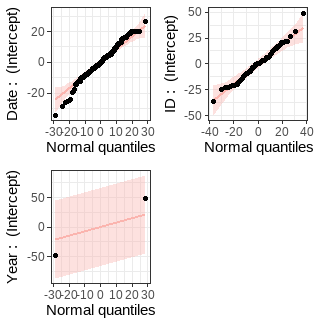

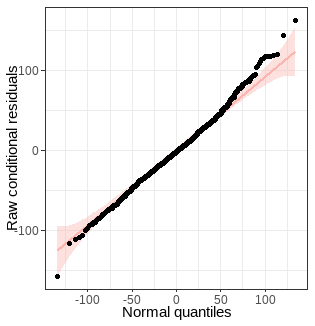
plots examined for significant deviation from reported expectations.

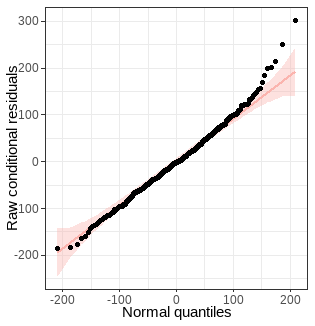

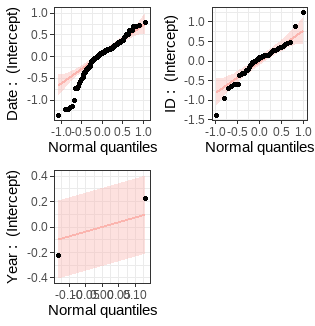

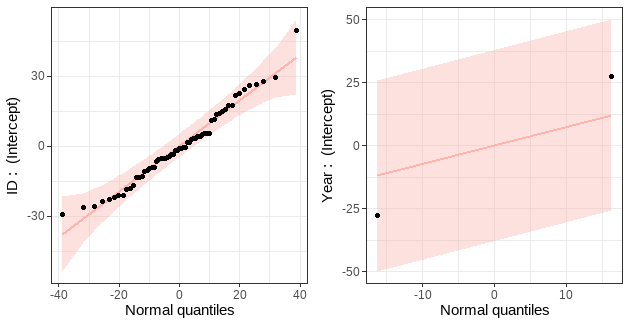
Supplementary Figure 2. Model: Individual Duration (Ind off-bout dur ~ 2 hr Temp)

Supplementary Figure 3. Model: Total Duration (24hr Constancy ~ 24hr Temp)

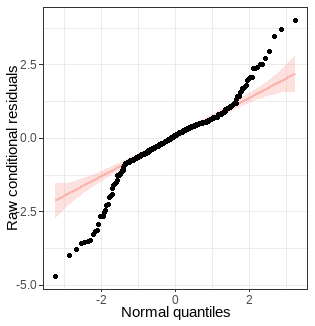

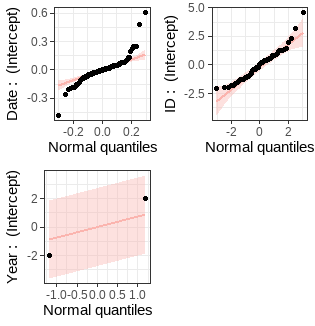

Supplementary Figure 4. Model: Morning off-bout start time. Extreme deviation from normality of our residuals suggests that this model does not meet requirements for model testing.
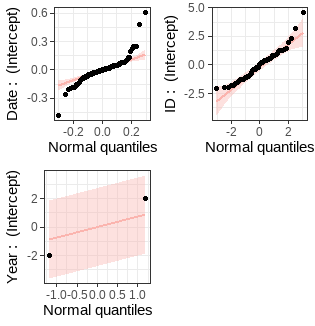

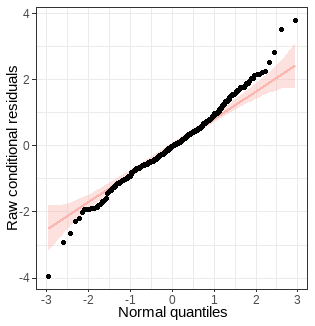

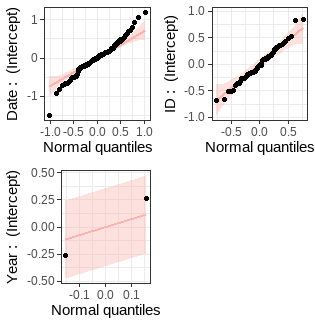

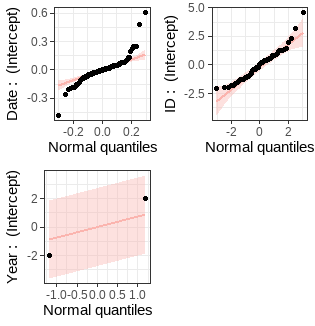

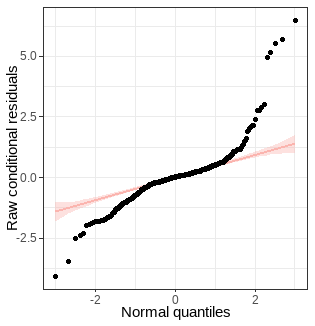
Supplementary Figure 5. Conditional residuals for model Afternoon off-bout start time. Extreme deviation from normality of our residuals suggests that this model does not meet requirements for model testing.

Supplementary Figure 6. Conditional residuals for model Total morning start time.
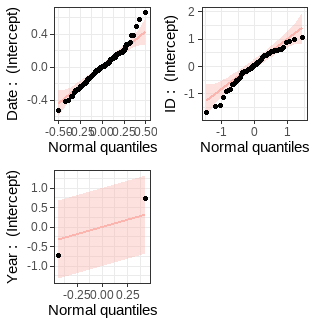

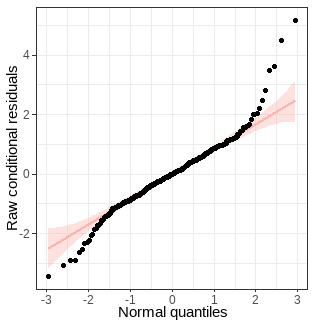

Supplementary Figure 7. Conditional residuals for model Total afternoon start time. Extreme deviation from normality of our residuals suggests that this model does not meet requirements for model testing.
